# LiF-MS+, a revised technique for mapping peptide-protein interactions

**DOI:** 10.1101/2024.05.28.596279

**Authors:** Benjamin Parker, Eric Weiss

## Abstract

Short linear motifs are sequences of amino acids present in unstructured polypeptide regions that function as ligands for specific sites on folded protein domains. These interactions, which often occur with low to modest affinity, modulate dynamic biological processes such as signal transduction and membrane trafficking. We recently described Ligand Footprinting-Mass Spectrometry (LiF-MS), a technique that rapidly and precisely maps sites at which short peptide ligands bind their biologically relevant recognition sites on folded protein domains. This approach marks the binding location of a peptide ligand on a structured protein using a cleavable crosslinker appended to the ligand that leaves behind a stable chemical modification following cleavage. This modification serves as a mass tag detectable by mass spectrometry, pinpointing sites of peptide ligand binding. Here we present LiF-MS+, an improved version of the footprinting technique that replaces the butanol mass tag with 1-butylpyrrolidine, which is positively charged at neutral pH and thus aids in ionization of the crosslinked peptide for analysis by mass spectrometry. We show ligand-mediated butylpyrrolidine footprinting effectively maps the well characterized binding interaction of the p38α mitogen-activated protein kinase (MAPK) with a MKK6 D-motif short linear motif peptide ligand, uncovering additional binding site information not observed in our original experiment. LiF-MS+ is thus a straightforward improvement of our previously published methodology for mapping the binding of short linear motifs to folded protein domains.

## INTRODUCTION

*Ab initio* computational prediction of peptide ligand binding to folded domains has high uncertainty, and crystallographic analysis of complexes of peptides and folded proteins is expensive, slow, and laborious. Additionally, crystallography often fails to capture structures with multiple dynamic conformations, and portions of co-crystallized peptide ligands may be invisible or not modeled into the electron density as is the case in multiple D-motif MAPK structures(1, 2). We developed Ligand Footprinting-Mass Spectrometry (LiF-MS), an approach that chemically maps sites of peptide ligand binding, to provide fine structure information about short linear motif – folded domain binding to inform computational prediction of structures and other experimental design.

LiF-MS exploits peptide ligands that carry a diazirine moiety that rapidly covalently crosslinks with nearby protein surfaces upon illumination with UV light. This reaction is relatively nonspecific, and does not require chemical groups such as primary amines or sulfhydryls on the target protein. Importantly, in LiF-MS the diazirine crosslinker is appended to a biotinylated peptide ligand of interest by an acid-labile linkage. Cleavage of this linkage eliminates the biotinylated peptide while leaving behind butanol moieties covalently attached at sites of crosslinking, providing mass tags that indicate likely sites of peptide ligand binding. This approach provides a way to discover novel peptide ligand docking sites and fill in poorly characterized regions in existing protein-peptide structures without more labor-intensive structural biology techniques.

In this work, we present LiF-MS+ (LiF-MS-Plus), a modified LiF-MS procedure that leaves a 125-Dalton 1–butylpyrrolidine mass tag following crosslinker cleavage, in place of the previously described 72-Dalton butanol moiety. Unlike butanol, butylpyrrolidine is positively charged at neutral pH. This assists in the ionization of the tryptic peptide during mass spectrometry, reducing representation biases caused by labeled peptides that ionize poorly. Here we show that LiF-MS+ produces a chemically defined mass tag which reliably maps the known peptide binding site of MKK6’s D-motif to the MAPK p38α. We validate the resulting site using the Trans-Proteomic Pipeline’s PTMProphet utility and demonstrate highly ranked crosslinked residues can be used as constraints for *de novo* docking site discovery.

## RESULTS

### LiF-MS+ places positively charged mass tags at peptide ligand interaction sites

Previously, we showed that LiF-MS can map the known docking site of D-motif peptide ligands to MAP kinases. However, as noted in that analysis, we found the butanol mass tag at precisely +72Da inconsistently, necessitating search for other chemically related tags with close masses such as butylamine (3) (see Supporting Information). Furthermore, crosslinker cleavage using previously established conditions (55C, pH 1.0 (4)) sometimes failed to produce mass tagged tryptic peptides representing the known D-motif. These inconsistencies could arise from processes that occur during linker cleavage, such as protein insolubility with resulting poor trypsin digestion and/or inefficient cleavage of the crosslinker. Alternatively, the butanol modification itself may suppress peptide abundance, such as by introducing excessive hydrophobicity during the LC step or through gas phase reactivity of butanol groups in MS-MS stages.

We therefore sought to replace the butanol mass tag with a positively charged adduct that increases solubility of labeled peptides and aids their ionization. We opted to cleave the crosslinker with pyrrolidine in the presence of catalytic formic acid (Figure 1A, B). This SN2 reaction leaves butylpyrrolidine, which is positively charged below pH ∼12 (Figure 1A, lower product), and a sulfamate leaving group. Importantly, pyrrolidine is a strong protein precipitant. Thus, to ensure efficient cleavage it is essential to denature diazirine crosslinked peptide-protein complexes in guanidine hydrochloride before pyrrolidine / formic acid treatment (see Materials and Methods). After crosslinker cleavage reaction the target protein is subjected to SDS-PAGE and in-gel trypsinization (Figure 1A; see bolded blue step), followed by LC-MS-MS analysis of the resulting proteolytic fragments. As discussed further below, we found that peptides with butylpyrrolidine modified residues were readily detectable by LC-MS-MS.

**Figure 1.**
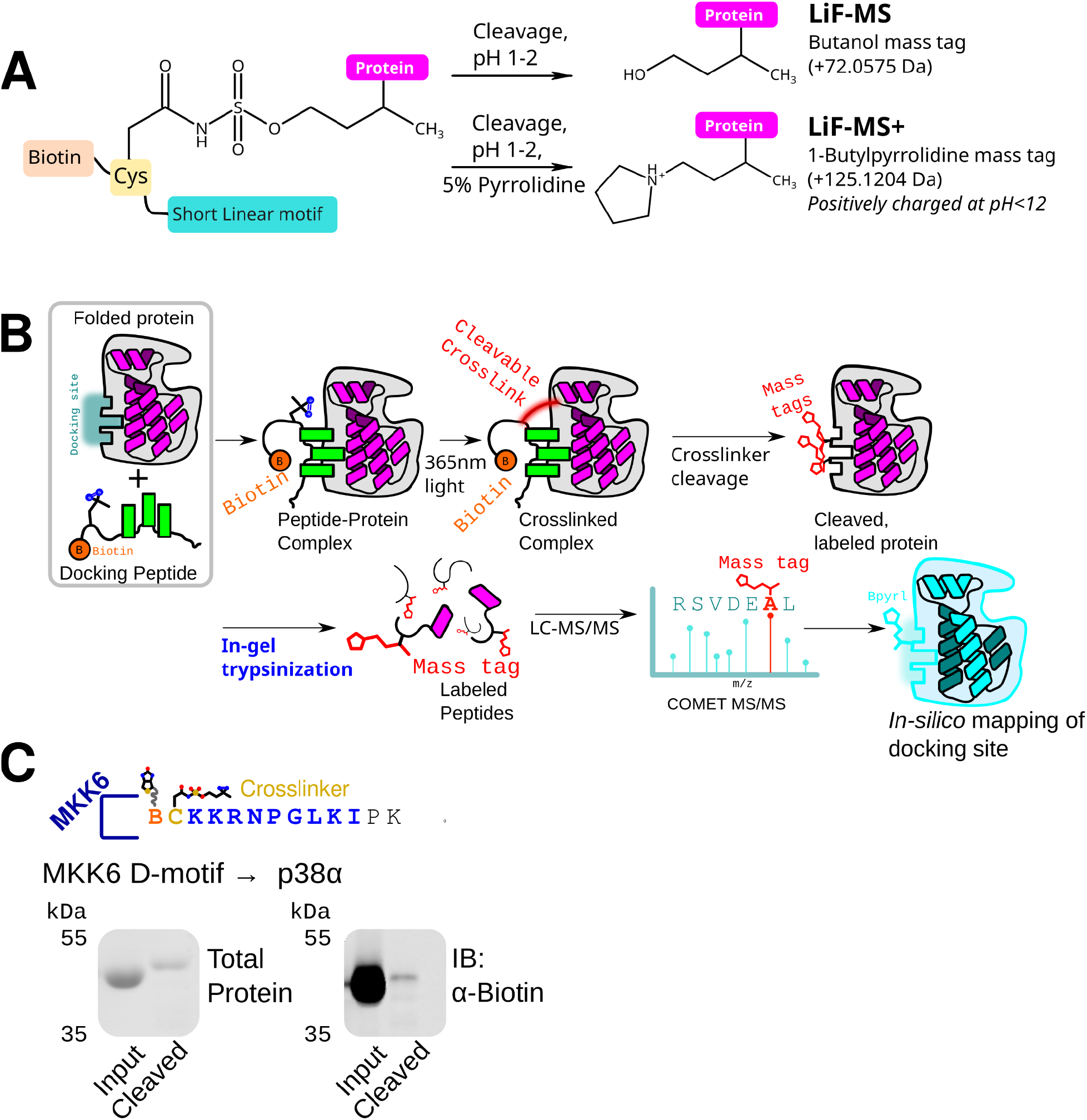
Overview and workflow of LiF-MS+, a revision of LiF-MS. **A**. Overview of LiF-MS+, showing mass tags and method of detection. **B**. Cleavage of the protein-peptide crosslinker using pyrrolidine yields the 1-butylpyrrolidine mass tag with a shift of 125Da instead of a butanol. **C**. Crosslinking and cleavage of the MKK6 D-motif to the MAPK p38α. Biotinylated peptide representing the MKK6 D-motif was crosslinked to purified p38α, followed by cleaving using the guanidine method (Materials and Methods). Left, total protein after crosslink as assayed by Revert™ Total protein stain (LI-COR Biotech). Right, blot probed IRDye® 800CW streptavidin before (Input) and after (Cleaved) cleavage of the crosslinker with pyrrolidine using the guanidine method (see Materials and Methods).

### LiF-MS+ maps the previously characterized MKK6-p38α docking interaction

In our description of the original LiF-MS approach we showed that it successfully maps the crystallographically and biochemically characterized site at which the MKK6 D-motif docking peptide ligand binds the MAPK p38α (1, 3). We therefore used the MKK6-p38α interaction to comparatively validate the LiF-MS+ approach. After performing diazirine crosslinking of the MKK6 peptide with p38α exactly as published previously (3) we cleaved the crosslinker with pyrrolidine and formic acid (see Materials and Methods). As expected for effective crosslinker cleavage, which should eliminate the biotinylated MKK6 peptide, pyrrolidine/formic acid treatment significantly reduced biotin present on p38α as shown by streptavidin western blot (Figure 1C).

We performed SDS-PAGE and in-gel trypsinization of the pyrrolidine/formic acid cleaved p38α protein, and then used the resulting gel-eluted peptides for LC-MS-MS analysis. Using COMET (5, 6), we searched for positions carrying the 1-butylpyrrolidine mass tag (+125.1204Da) (see Materials and Methods). We scored the location of butylpyrrolidine modified residues using the confidence score derived from the reciprocal of the COMET E-value, as described for the original LiF-MS technique (3). As shown in figure 2A, residues 122-126 scored as crosslinking sites with highest confidence, with residues 161-165 scored as crosslink sites with lower confidence. A third region, 299-304, had crosslink confidence scores about an order of magnitude lower. We also searched for high confidence butylpyrrolidine modified residues using the PTMProphet utility from the Trans-Proteomic Pipeline (7, 8). This identified residues 123 and 161 as likely crosslink sites, in excellent agreement with COMET analysis (Figure 2B).

**Figure 2.**
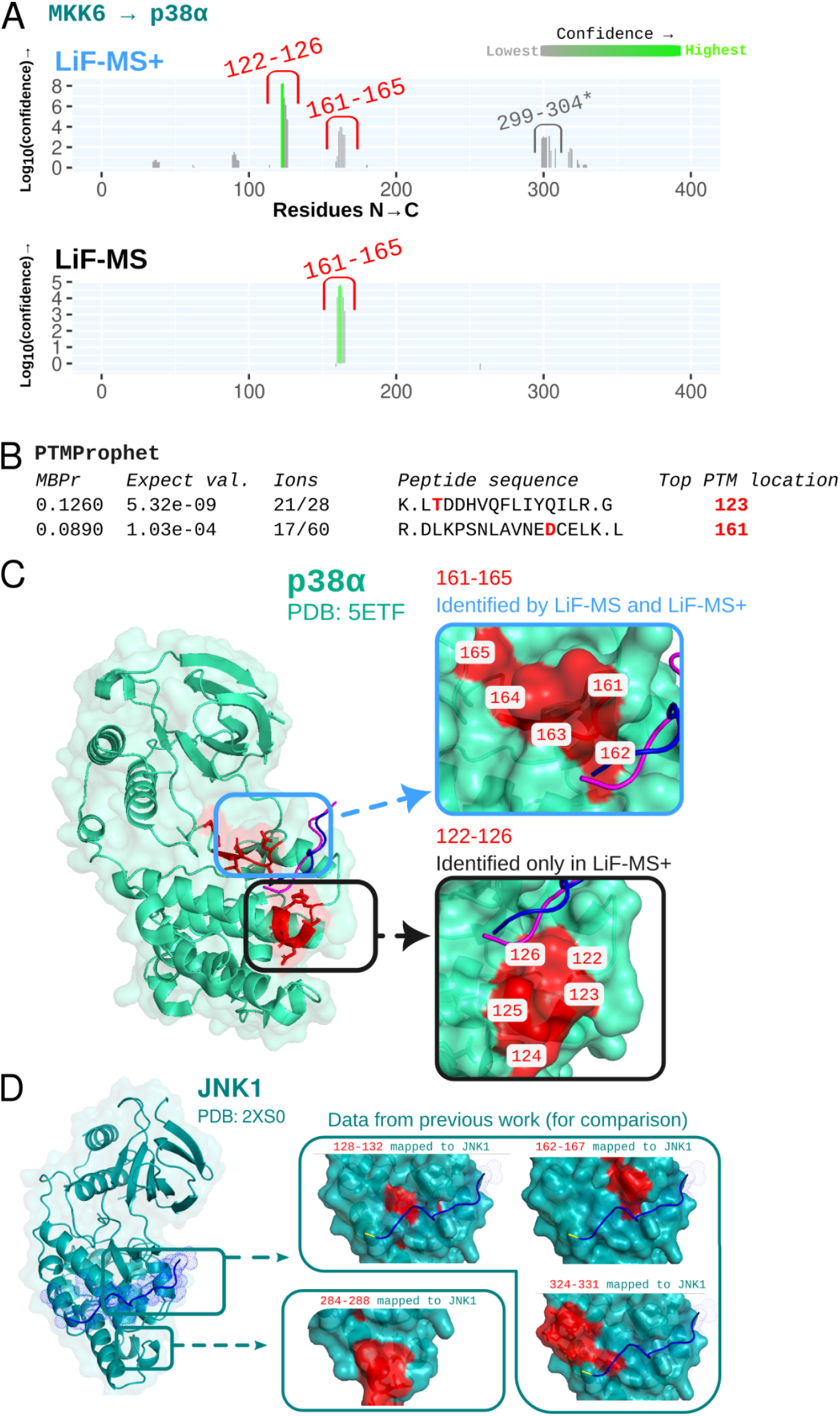
LiF-MS+ identifies the binding site of the MKK6 D-motif on p38α. **A**. Confidence scoring of residues modified with 1-butylpyrrolidine (+125.1204Da). Residues consistent with the published docking site are highlighted in red. Grey residues marked with * are not close to the docking site and are likely an off-target binding or crosslinking event. **B**. Top two peptides with 1-butylpyrrolidine tags as assayed by the PTMProphet utility in the TPP pipeline, using a cutoff of 0.001 for both probability (MBPr) and Expect value. **C**. Crosslinked amino acid residues from A and B mapped onto the p38α structure (PDB: 5ETF) shown in red. D-motif peptides from the p38α-MKK6 structures 5ETF and 2Y8O are shown in blue and purple, respectively. Crosslinks identified by both LiF-MS+ and LiF-MS are highlighted in blue; crosslinks identified only by LiF-MS+ are highlighted in black. **D**. LiF-MS analysis of MKK4 D-motif binding with Jnk1 from our previous work showing two distinct crosslink sites closely resembling the LiF-MS+ data for p38α.

As shown in Figure 2C, the highest scoring butylpyrrolidine modified residues closely align with the N-terminus of the MKK6 D-motif from published crystal structures of the MKK6-p38α complex (1, 2). One of the regions, residues 161-165, was identified in prior LiF-MS analysis of the complex (3)(Figure 2B, blue outline). However, LiF-MS+ alone strongly identified crosslinking of the 122-126 region (Figure 2B, black outline). Notably, both of these crosslink sites closely reflect our prior LiF-MS analysis of the MKK4-JNK1 docking interaction (3) (Figure 2D).

### Butylpyrrolidine tagged residues can be used as constraints for docking cleft discovery

We next determined if we could recapitulate the p38α docking site using constraints generated from LiF-MS+. We used the CABSdock utility(9) and the workflow we published previously(3), constraining the N-terminus of the MKK6 D-motif to the highest ranked crosslinked residues 122 and 123 (Figure 3A). The confidence scores of these residues is two orders of magnitude higher than the next set of crosslinked residues (Figure 2A). We merged the output peptide structures and filtered out structures not within 8Å of the indicated residues. We then hierarchically clustered the structures by RMSD and chose the largest cluster, displayed in Figure 2B. Averaging the backbone coordinates of the largest cluster of CABSdock structures, we found the putative CABSdock solution closely matched the reference MKK6 D-motif structure (Figure 3C).

**Figure 3.**
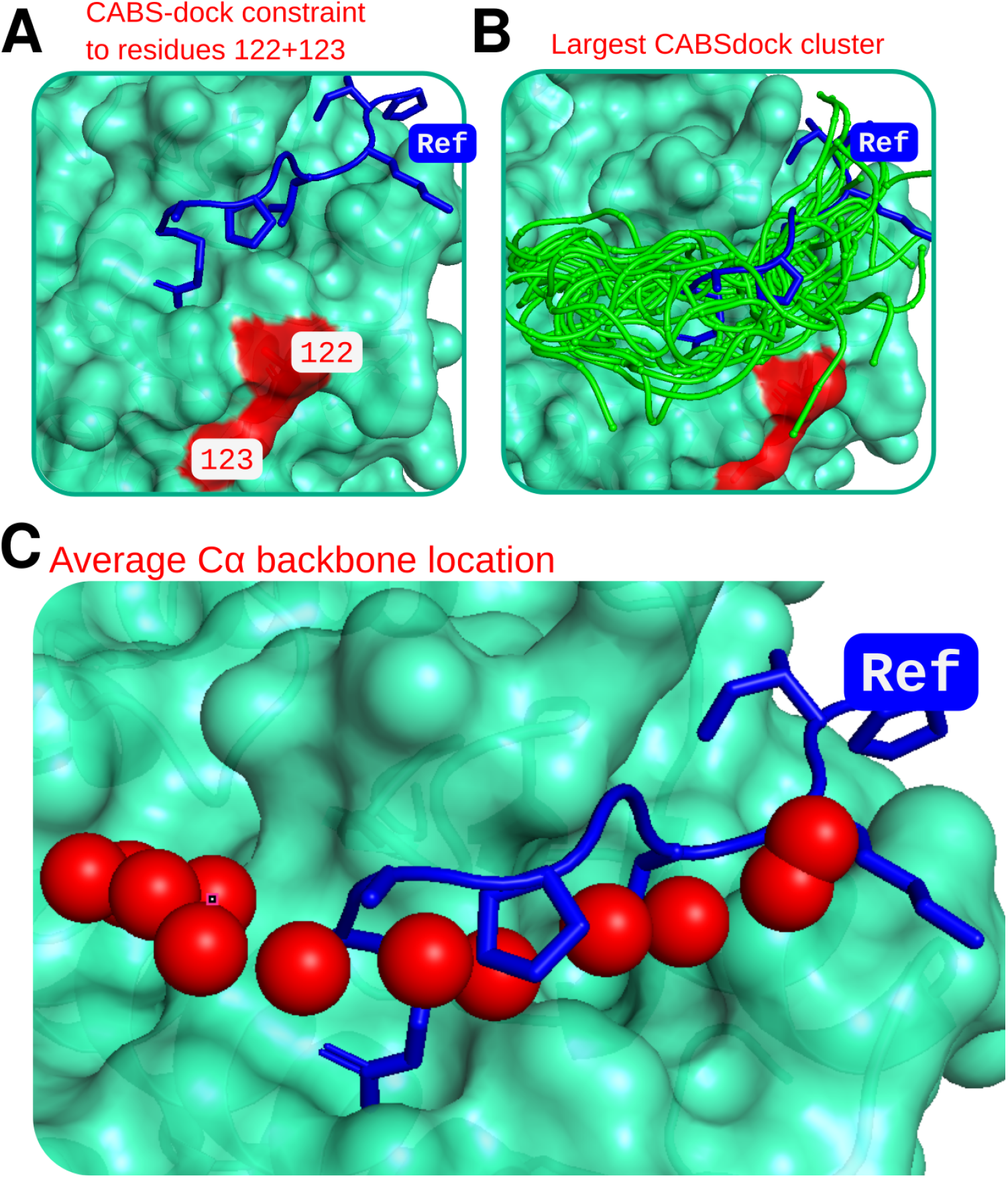
Residues tagged with 1-butylpyrrolidine can be used as constraints for CABSdock-mediated docking site discovery. **A**. Residues 122+123 were used as constraints for CABSdock modelling, as these were two orders of magnitude higher confidence than other residues. **B**. The largest of 10 CABSdock clusters after filtering out peptide models not within 8 angstroms of the indicated residues. **C**. Average backbone Cα location for peptide models in the cluster shown in B, showing alignment with the reference peptide from the crystal structure. In all cases, the crystal structure of the MKK6-p38α complex is shown (PDB: 5ETF), with the kinase shown in green and the MKK6 D-motif shown in blue sticks (labelled “Ref”).

## DISCUSSION

The LiF-MS methodology we originally described is clearly able to identify sites of peptide ligand binding on folded protein domains. However, unpublished data from our group suggests that the protocol’s reliance on hydrophobic butanol mass tags might suppress the abundance of some tryptic peptides in LC-MS-MS analysis. Potential effects of butanol mass tags could potentially depend strongly on peptide context, reducing the abundance of some tryptic fragments while exerting little influence on others. Notably, we found that the positively charged mass tag butylamine, a potential minor breakdown product from crosslinker cleavage, could be used instead of butanol in some cases to map peptide docking sites (3) (See Supplementary Methods). We therefore developed the LiF-MS+ protocol that uses pyrrolidine / formic acid cleavage as a chemically defined method to create a positively charged mass tag at crosslink sites in higher abundance, which likely boosts ionization of tagged peptides.

Our LiF-MS+ analysis of the MKK6-p38α complex indicates that butylpyrrolidine is an effective crosslinker-directed mass tag that clearly identifies the well characterized MKK6 D-motif ligand binding site on p38α. Prior LiF-MS analysis identified only the p38α surface comprising residues 161-165 as close to the D-motif peptide N terminus. In contrast, LiF-MS+ identifies two regions as near the D-motif N terminus: residues 122-126 and, with lower confidence, residues 161-165. As noted, these newer data for the MKK6-p38α complex are more consistent with LiF-MS analysis of MKK4-JNK1, suggesting that the butylpyrrolidine mass tag may provide a more complete picture of ligand-directed crosslink sites.

### Technical considerations

We note that the tryptic peptide crosslinked at residues 161-165 was not found in the mass spectrometry data unless COMET was set to not search for iodoacetamide (IAA)-alkylated (+57Da) cysteines. Indeed, this peptide contains a cysteine at position 162, which is within the p38α docking site (Figure 2C). This suggests that residues modified with 1-butylpyrrolidine within 5 or so amino acids may block efficient alkylation during sample preparation. While we did not try removing IAA from the preparation process, we suggest searching for peptides both and without the alkylated cysteine requirement to ensure potential tryptic peptides aren’t missed.

It is important to score crosslinked amino acid residues relative to each other beyond a base probability level. We have analyzed our data with the COMET utility(5, 6), PTMProphet(8), as well as the FragPipe utility and found COMET provides the greatest degree of resolution of the docking site, as each crosslinked tryptic peptide discovered has an associated raw E-value which can be used to calculate our confidence score. Notably, while FragPipe discovers high-confidence butylpyrroldine-modified sites, it scores on target and off target sites roughly equally, and thus is not useful for this application.

## Materials and Methods

### Purification and crosslinking of MKK6-p38α

Purification and crosslinking was performed as described (3). Briefly, we incubated ∼50 ug His-tagged p38α in 10 ul crosslinking buffer (50mM Tris pH 8.3, 150mM NaCl) containing ∼35 uM peptide at 4C and irradiated with a 365nm UV lamp for 20 minutes.

### Cleavage and butylpyrrolidine formation

We added 10 volumes (100 microliters) of 8M guanidine hydrochloride to one volume of solution containing the crosslinked protein-peptide complex to denature the protein. We then added, in order, formic acid to 5% (v/v) and pyrrolidine to 5% (v/v). After heating at 55 degrees for 2 hours to cleave the crosslinker, we diluted the solution 1:3 with water and added 4 volumes of acetone (post dilution) to precipitate the protein. We then redissolved the pelleted protein in 20μL 7M urea, 2M thiourea, 1% SDS, and 1M ammonium bicarbonate without boiling. The resulting solution was run on 12% SDS-PAGE, stained with SimplyBlue™ SafeStain (Invitrogen), and gel extracted.

### Mass spectrometry and scoring

We performed LC-MS-MS, scoring, and CABSdock/docking site discovery as described (3). For this analysis we searched for 1-butylpyrrolidine missing two hydrogens (125.1204 Da).

Using high-confidence modified residues 122 and 123 of p38α we performed CABSdock/docking site discovery as described (3). We performed one CABSdock run per crosslinked residue with a 14Å constraint to the MKK6 D-motif N-terminus. We removed modeled peptide structures with backbone atoms further than 8Å from the site. We hierarchically clustered the resulting structures by RMSD into 5 clusters per crosslinked site (10 clusters total). Figure 3 shows the largest cluster.

Mass spectrometry data can be downloaded at the following link: https://drive.google.com/drive/folders/1CSXnO1ePSVLYCFkKs7lFFkOUOrAMTtTI?usp=sharing

## Acknowledgements

Proteomics services were performed by the Northwestern Proteomics Core Facility, generously supported by NCI CCSG P30 CA060553 awarded to the Robert H Lurie Comprehensive Cancer Center, instrumentation award (S10OD025194) from NIH Office of the Director, and the National Resource for Translational and Developmental Proteomics supported by P41 GM108569.

